# Structural basis of SALM3 dimerization and synaptic adhesion complex formation with PTPσ

**DOI:** 10.1101/2020.01.09.893701

**Authors:** Sudeep Karki, Alexander V. Shkumatov, Sungwon Bae, Jaewon Ko, Tommi Kajander

## Abstract

Synaptic adhesion molecules play an important role in the formation, maintenance and refinement of neuronal connectivity. Recently, several leucine rich repeat (LRR) domain containing neuronal adhesion molecules have been characterized including netrin G-ligands, SLITRKs and the synaptic adhesion-like molecules (SALMs). Dysregulation of these adhesion molecules have been genetically and functionally linked to various neurological disorders. Here we investigated the molecular structure and mechanism of ligand interactions for the postsynaptic SALM3 adhesion protein with its presynaptic ligand, receptor protein tyrosine phosphatase σ (PTPσ). We solved the crystal structure of the dimerized leucine rich repeat (LRR) domain of SALM3, revealing the conserved structural features and mechanism of dimerization. Furthermore, we determined the complex structure of SALM3 with PTPσ using small angle X-ray scattering, revealing a 2:2 complex similar to that observed for SALM5. Solution studies unraveled additional flexibility for the complex structure, but validated the uniform mode of action for SALM3 and SALM5 to promote synapse formation. The relevance of the key interface residues was further confirmed by mutational analysis with cellular binding assays and artificial synapse formation assays. Collectively, our results suggest that SALM3 dimerization is a pre-requisite for the SALM3-PTPσ complex to exert synaptogenic activity.

## Introduction

Neurons form highly complex networks of specific connection patterns through the cellular junctions at synapses. Synaptic adhesion molecules are emerging synapse organizers, which are localized at the nerve cell membranes at the synaptic cleft, and play a crucial role in the formation, maintenance and refinement of neural circuits^1^. Several families of synaptic adhesion molecules have been identified, including e.g. neurexins^2^, neuroligins^3^, synaptic adhesion like molecules (SALMs)^4^, leucine-rich repeat (LRR) transmembrane neuronal proteins (LRRTMs)^5^, leukocyte common antigen-related receptor protein tyrosine phosphatases (LAR-RPTPs)^6^ and netrin-G ligands (NGLs)^7^. The dysfunction of synaptic adhesion molecules have been proposed to be involved in onset and progression of various neurological disorders, which includes autism spectrum disorders, schizophrenia and Alzheimer’s disease^8-11^.

The SALM family of proteins also known as LRR and fibronectin III domain containing proteins (LRFNs), represent a family of LRR-containing synaptic adhesion molecules, with five known members (SALM1-SALM5)^4^. All the SALM proteins share a similar domain architecture consisting of an extracellular region with an LRR domain, an immunoglobulin (Ig) domain and a fibronectin III (FnIII) domain followed by a transmembrane domain (TM) and a non-structured cytoplasmic region^12^. Notably, SALM1-3 contain type I PDZ binding motifs, which are absent in SALM4 and SALM5^4, 12^ (Fig. 1a).

**Figure 1.**
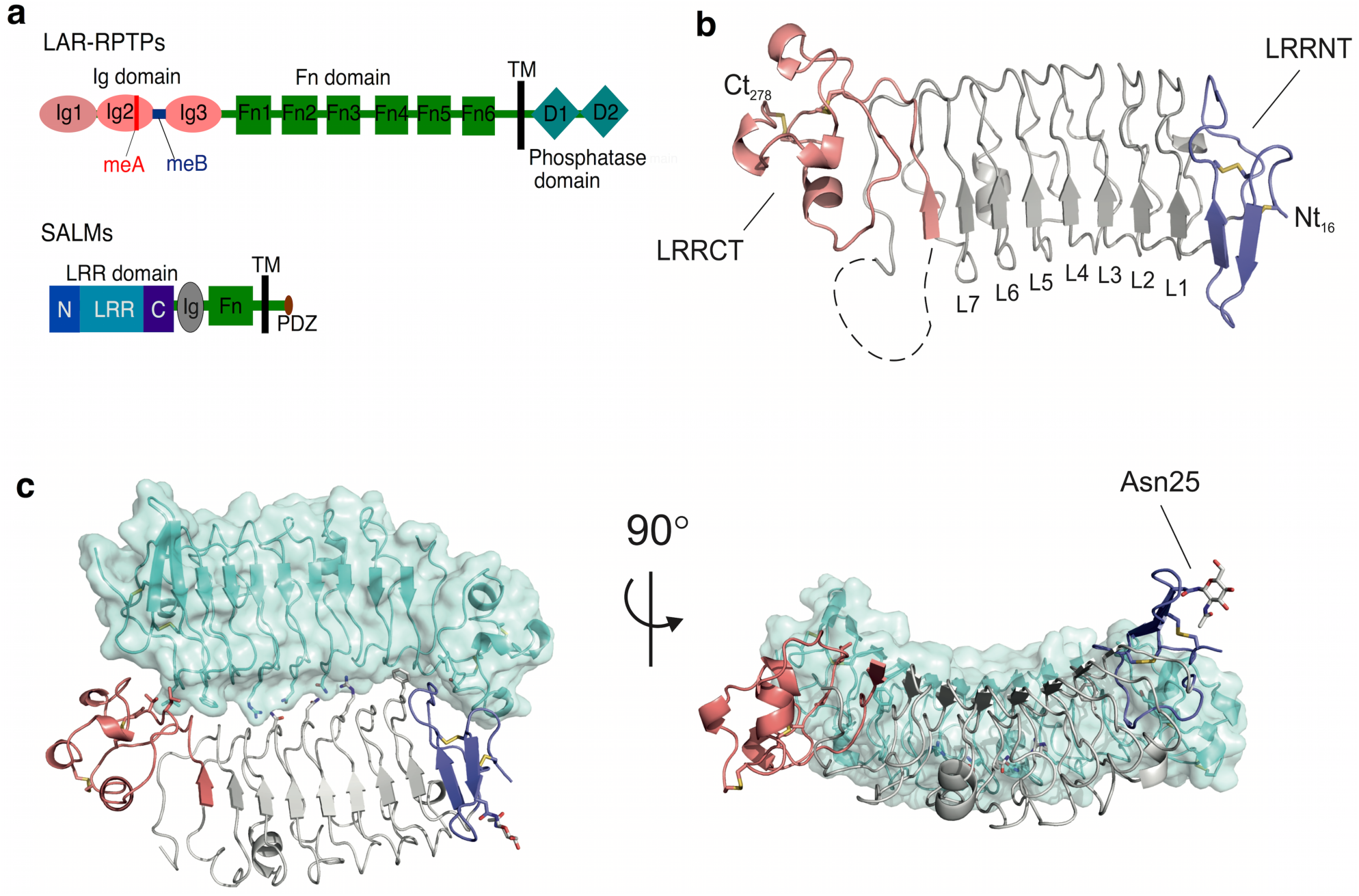
Crystal structure and dimer assembly of SALM3 LRR domain. a) Domain organization of SALMs and LAR-RPTPs. Main features are indicated as: Ig, immunoglobulin domain; Fn, fibronectin domain; Tm, a transmembrane domain; D, phosphatase domain; LRR, LRR domain; N, LRRNT; C, LRRCT; PDZ, PDZ binding motifs. b) The overall SALM3 LRR domain structure. The missing parts for the LRR domain shown as dashed lines, LRRNT and LRRCT-capping domains are colored blue (LRRNT) and red (LRRCT) respectively. The N- and C-termini of the protein are marked with ‘Nt’ and ‘Ct’ and the residue number, disulphides are indicated in yellow as sticks. c) Dimeric assembly observed in the crystal structure with one of the subunits shown with transparent surface, and N-glycosylation present in the LRRNT Asn25 shown as sticks. On the left structure is viewed from the “top” towards the concave side of the LRR dimer, orthogonal to the interface, and on the right from the side, glycosylated Asn25 marked.

SALMs are considered to be postsynaptic adhesion proteins, mostly expressed in the neurons and regulate neurite outgrowth and branching^12^. In addition, they have been implicated in the regulation and development of synapse formation^12-14^. SALM3 and SALM5 possess synaptogenic activity, and promote both excitatory and inhibitory presynaptic differentiation in contacting axons^15^ via *trans*-synaptic interaction with LAR-RPTPs^13, 14^. SALM4 suppresses excitatory synapse development by *cis*-inhibiting *trans*-synaptic SALM3-LAR-RPTP adhesion^16^. Crystal structures of human SALM5-PTPδ, and also SALM2-PTPδ complexes reveal 2:2 heterotetrameric *trans*-synaptic association between the proteins^17, 18^. We previously reported the crystal structure of mouse SALM5 LRR-Ig domain fragment, and solution structures of SALM3 and SALM5 extracellular regions^19^. This study revealed that LRR domains of SALM5 are involved in dimerization, and the different structures indicated that overall structural features are conserved between SALM2, SALM3 and SALM5^19^. However, structural details of SALM family beyond SALM2 and SALM5 remain to be determined including SALM3 and the molecular details of its interaction with LAR-RPTPs.

SALM3 is expressed early during embryonic stage in the brain, whereas the other SALMs are expressed from the embryonic stage, and the expression is maintained or increased after birth^20^. *In vivo* studies using SALM3 knockout (SALM3^-/-^) mice displayed markedly reduced excitatory synapse number and locomotor activity, and exhibit behavioral hypoactivity, which underlines the importance of SALM3 in excitatory synapse development^14^. SALM3 interacts with all three known members of LAR-RPTPs (LAR, PTPσ and PTPδ) to induce presynaptic differentiation^14^. The interaction between SALM3 and LAR-RPTPs is markedly enhanced in the presence of the LAR-RPTP mini-exon B (meB)^14, 19^. Besides SALMs, LAR-RPTPs form *trans*-synaptic complexes with different families of postsynaptic adhesion proteins, which include NGL-3, IL1RAcP, neurotrophin receptor TrkC and the Slitrks^21^. The extracellular region of LAR-RPTPs contain three Ig domains, four to eight FnIII domains and multiple splicing variants at several sites: mini-exon A (meA) in the Ig2 domain, mini-exon B (meB) between Ig2 and Ig3 domain and mini-exon C (meC) located in the FnIII domain (**Fig. 1a**). In the cytoplasmic region, LAR-RPTPs contain two tandem tyrosine phosphatase domains^6, 21^. The Ig2 and Ig3 domains of PTPδ and PTPσ are involved in interaction with the LRR and Ig domains of SALM5^17, 18^.

In this study, we have solved the crystal structure of mouse SALM3 LRR domain and show that it forms a dimer that is close to identical to that of SALM5, and the residues that are involved in dimerization and their interactions are largely conserved, and our complementary biophysical studies support this. We also solved the small angle X-ray scattering (SAXS) structure of SALM3-PTPσ complex that verifies the formation of the 2:2 *trans-*heterotetrametric complex between the proteins. Mutational studies with a cell-based binding assay and a synapse formation assay confirm the binding mechanism of SALM3 to PTPσ and the identity of the key interface residues that are largely the same as for the SALM5-PTPδ interaction. Therefore, we conclude that SALM3 and SALM5 interact in similar manner with their LAR-RPTP ligands to induce presynaptic differentiation. Solution scattering studies suggest that the dimeric complex has significant conformational flexibility. Our results also confirm that dimerization is absolute requirement for induction of synapse formation by the SALM3, similar to SALM5, and thus appears to be common for the SALM family proteins involved in synapse formation.

## Results and Discussion

### The structure of the SALM3 LRR domain

The structure of mouse SALM3 LRR domain (**Fig. 1b, c**) was solved at 2.8 Å resolution in space group P2_1_ **(Table 1)**. The crystals exhibited high degree of twinning, making the structure solution demanding (see methods). However, it was possible to obtain a good quality refined model once the correct space group was resolved. Four molecules, or two dimers, were present in the asymmetric unit of the crystal, one monomer had an N-glycan partly visible in electron density at Asn25 at the N-terminal end of the domain (**Fig. 1c**). Overall two N-glycosylation sites are predicted to be present in SALM3 LRR domain (Asn25 and Asn70), with most of the glycans disordered in the structure, however. The LRR domain shows a typical extracellular LRR domain structure, similar to the known SALM5 LRR structure (Supplementary **Fig. S1)**^19^, with an extend β-sheet on the concave side of the structure formed by the seven LRRs (L1-L7) (**Fig. 1b**) of variable length (22-24 residues) with consensus motif of “L_1_xxL_4_xL_6_xL_8_xxN_11_xL_13_” with aliphatic Leu or Ile residues mostly occupying positions 1, 4, 5, 6 and 13 and forming the hydrophobic core of each repeat, and the absolutely conserved Asn11 hydrogen bonding to the previous repeat. The LRR domain further contains an N-terminal LRRNT capping subdomain (residues 17-48) and an α-helical C-terminal LRRCT capping domain (residues 228-278) **(Fig. 1b**,**c)**, which are conserved in the vertebrate extracellular LRR domain family^22^. Both the LRRNT and LRRCT subdomains of SALM3 are stabilized by two disulphide bridges; in the LRRNT between Cys17-Cys27 and Cys21-Cys34, and in the LRRCT between Cys238-Cys257 and Cys240-Cys278. Cys278 also is the last residue of the whole LRR domain. A flexible loop (residues 216-227) is present between the 7^th^ LRR and the LRRCT, not visible in the SALM3 crystal structure **(Fig. 1b, c**). The possible functional significance of this loop remains unknown, as its not involved in LAR-RPTPs recognition at least in SALM5-PTPδ complex structures^17, 18^, while such an extension might be expected to have some functional significance, as *e*.*g*. in the case of platelet glycoprotein IB alpha LRR domain^23^.

**Table 1.** Crystallographic data collection and refinement statistics.

| **Data collection parameters** |  |
| --- | --- |
| Beam line | DLS I03 |
| $\lambda$ (Å) | 0.97625 |
| Temperature (K) | 100 |
| Space group | P2<sub>1</sub> |
| Unit cell |  |
| a, b, c (Å) | 31.45, 132.16, 134.18 |
| $\beta$ (°) | 90.1 |
| Resolution (Å) | 50-2.8 |
| Observations | 158799 |
| Unique | 26991 |
| R<sub>merge</sub> (%)* | 17.5 (119) |
| CC(1/2)* | 99.5 (70) |
| I/ (I)$\sigma$* | 9.0 (2.1) |
| **Refinement** |  |
| Resolution (Å) | 50-2.8 |
| R<sub>work</sub>/R<sub>free</sub> (%) | 26.91/28.50 |
| No. of reflections /R<sub>free</sub>-set reflections | 26902/1347 |
| Average B-factor (Å<sup>2</sup>) | 88.27 |
| No. of atoms (protein) | 7258 |
| r.m.s.d bond lengths (Å) | 0.006 |
| r.m.s.d bond angles (°) | 0.88 |
| Ramachandran plot (favoured/outliers)(%) | 92.24/0.6 |

*\*Statistics for highest resolution shell are given in parentheses.*

### Dimerization through the LRR domain is a defining feature of the SALM family

The SALM3 LRR domain forms a dimer similar to those observed for SALM5 and SALM2 crystal structures^17-19^ (**Fig. 1**). It appears that the SALMs form a highly conserved dimeric structure based on the known structures of SALM2, SALM3 and SALM5 and our data on SALM1 oligomerization^19^, and the overall sequence conservation of the interface residues in the protein family (Supplementary **Fig S2, Table S1**). The SALM3 LRR structure and dimer interface are close to identical to that of SALM5 (**Fig. 2**, Supplementary **Fig S1, Table S1**)^19^. The hydrophobic interactions and hydrogen bonding networks are conserved at the interface with Gln131 and Arg107 forming hydrogen bonds across the dimer interface to Asn156 of the other monomer and *vice versa* (**Fig. 2, Table S1**). The SALM3 LRR domain, like SALM5 and SALM2 forms an antiparallel side-by-side dimer, with a rather small interface with buried surface area of 1001 Å^2^. The interface consists of the “one-layer” central hydrogen bonding network (**Fig. 2**) described above, and hydrophobic interactions formed by the N- and C-terminal regions (**Fig. 2**). The dimer species is however very stable in solution for both SALM3 and SALM5, and also SALM1 based on our observations^19^. Monomeric species have not been observed *in vitro*. Here, we analysed the oligomeric state in solution for both the LRR domain and LRR-Ig construct for SALM3 by size exclusion chromatography-coupled multiangle static laser light scattering (SEC-MALLS) (**Fig. 3b**,**c**). Both appear as dimers based on observed molecular weights of ca. 60 and 80 kDa (**Fig. 3b**,**c**); twice the size of molecules weights calculated from sequence (30,9 and 40 kDa). Thus, these measurements verified that the LRR domain is sufficient alone for stable dimer formation. Comparison of the SALM3, SALM5 as well as SALM2 dimers reveals highly similar structures with r.m.s.d. values of 0.691 and 0.758 Å for the LRR domains of SALM5 and SALM2 against SALM3 structure, respectively. Differences can be seen mainly in the conformation of the LRRNT capping region (Supplementary **Fig. S1**). Thus, we conclude that the dimerization mechanism through the LRR domains is conserved in the family, generating a unique dimeric post-synaptic LAR-RPTP ligand, with possible functional implications. Interestingly all the other characterized LAR-RPTP ligands (Slitrks, IL1RAPL1 and TrkC and NGL-3)^24-27^ are monomeric. Dimerization of SALMs might be a way of regulation of the function and induction of presynaptic signaling by LAR-RPTPs, as in the case of SALM3 and SALM5 biological function depends on the oligomerization^17, 18^ (see below), and SALM4 regulates the function of SALM3 by *cis*-inhibition, most likely through heterodimer formation^16^. Function of SALM2 remains unclear, but it appears to be able to bind the LAR-RPTP receptors^18^, whereas the function of SALM1 appears quite different, with reported involvement in actin mediated post-synaptic neurexin clustering, and no synapse formation activity reported ^28^.

**Figure 2.**
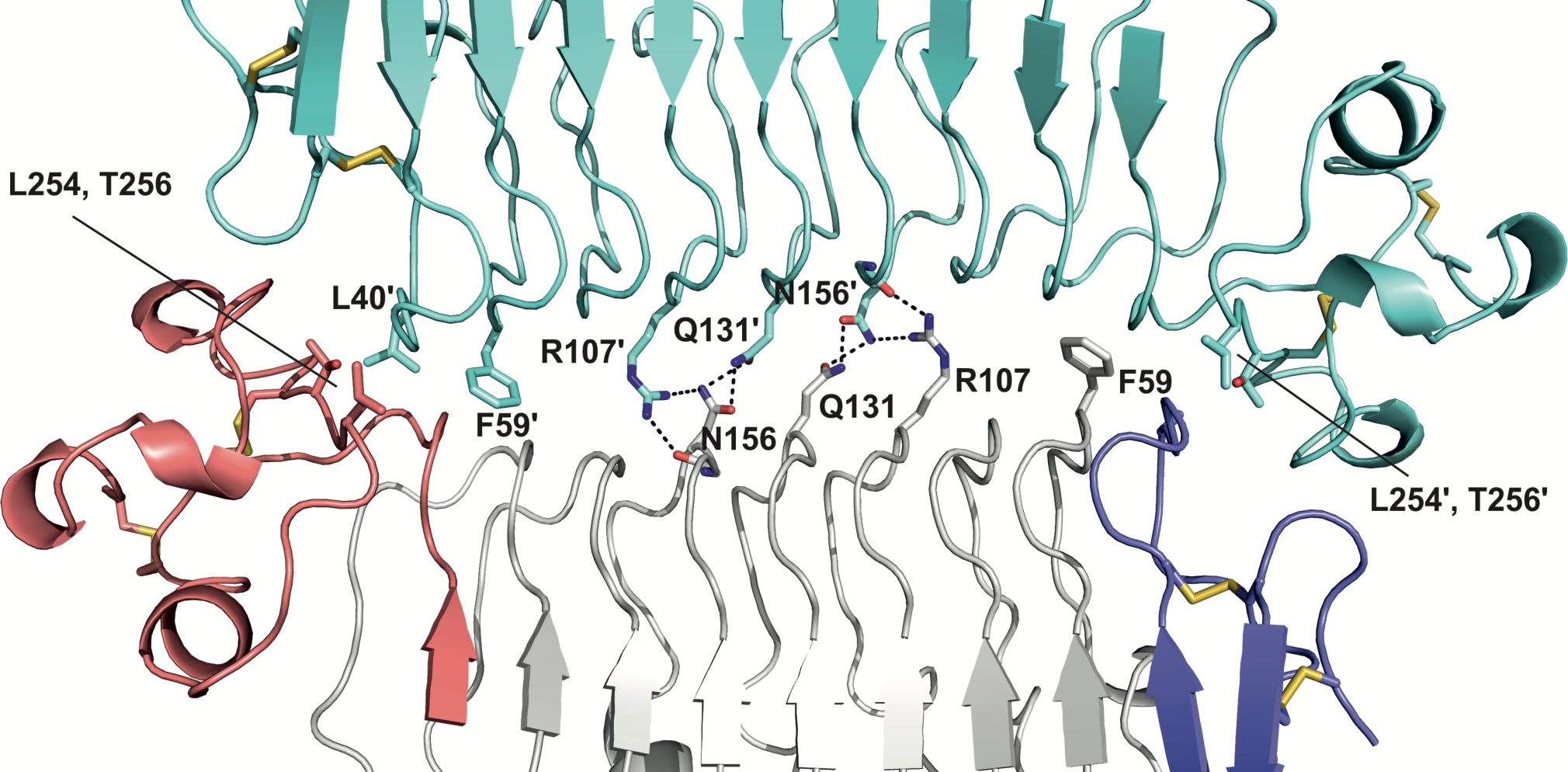
The dimer interface of SALM3 LRR domain. Residues contributing to the interface either by hydrogen bonds (dashed lines) or hydrophobic interactions are indicated. One monomer (top) is shown in cyan, and the other in grey with N-terminal capping LRRNT-subdomain in blue and LRRCT-subdomain in red, disulphides indicated in yellow as sticks.

**Figure 3.**
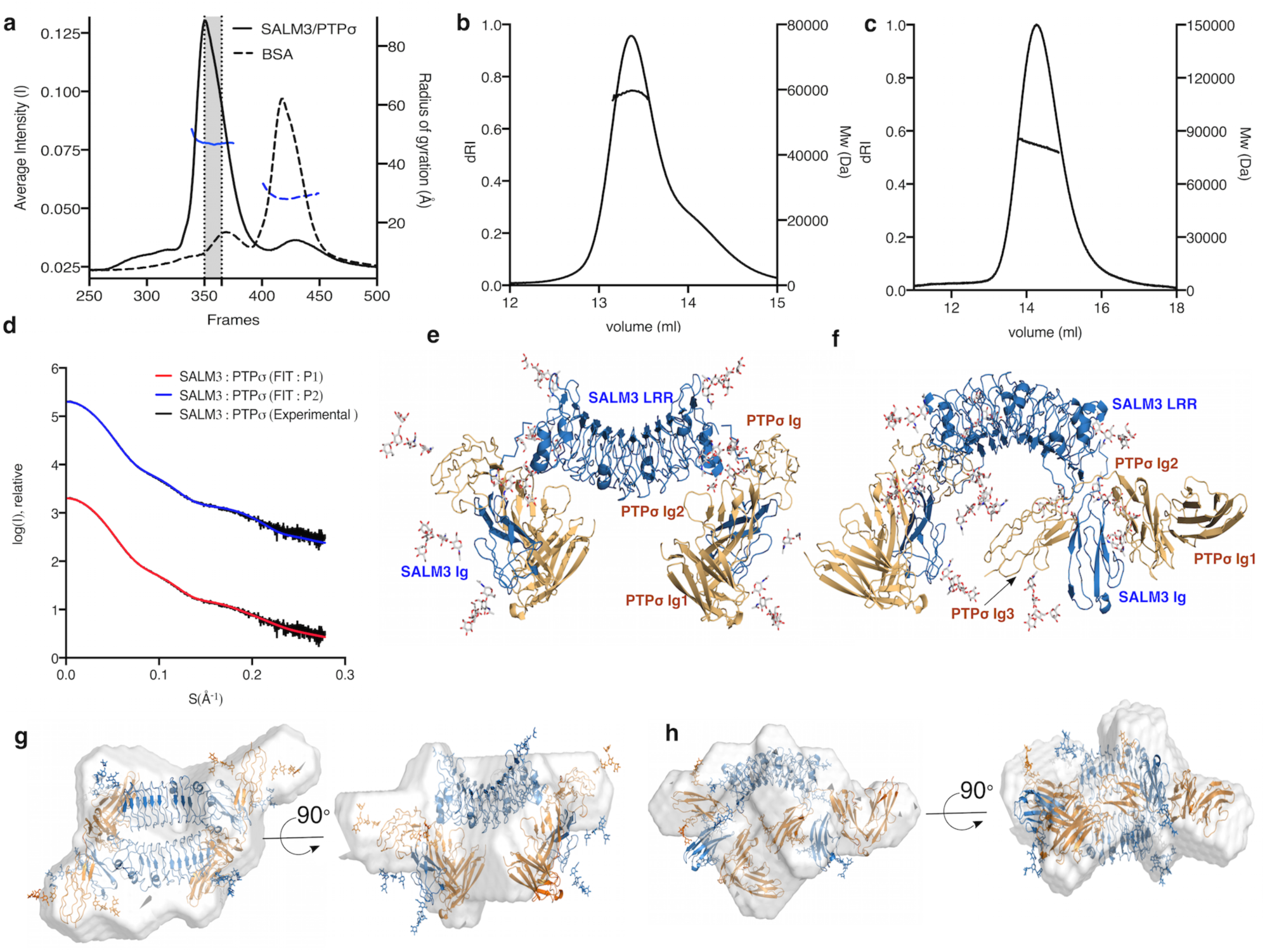
Solution structure of the complex between SALM3 and PTPσ. a) The SEC-SAXS profile of SALM3-PTPσ complex (solid line) and BSA (dashed line). Frames (350-365, shaded regions) were selected for further SAXS analysis. Rg-values (left Y-axis) were plotted over averaged X-ray scattering intensities (right Y-axis). b) and c) Oligomeric state of SALM3 LRR and SALM3 LRR-Ig constructs respectively, characterized by analytical size exclusion chromatography and static light scaterring. Calculated Mws (right Y-axis) are plotted over the protein (dRI) signal peaks; average observed molecular weights were 59.5 kDa and 82.4 kDa for LRR and LRR-Ig constructs, respectively. d) Scattering intensity profiles with the corresponding fit to SALM3-PTPσ complex models generated with rigid body modeling in CORAL in P1 and P2 symmetry. Experimental SAXS profiles were appropriately displaced along the logarithmic axis for better visualization and overlaid with corresponding fits. e) and f) show the SALM3-PTPσ complex models in P2 and P1 symmetry, respectively. SALM3 LRR and Ig domains are colored blue, PTPσ Ig1-3 domains are colored orange and modelled glycans are colored by atom (carbon: white, oxygen: red and nitrogen: blue). g) and h) Superimposition of the *ab initio* calculated envelope models (grey) and SALM3-PTPσ rigid body complex models with P2 and P1 symmetry, respectively.

### The solution structure of SALM3-PTPσ complex

We have characterized the solution structure of SALM3 in complex with its PTPσ receptor, in order to understand the binding mode to the ligand for SALM3 by SAXS. Size exclusion chromatography coupled with SAXS (SEC-SAXS) was used to obtain the SAXS data on the SALM3_LRR-Ig_-PTPσ_Ig1-Ig3_ complex to characterize the interaction between SALM3 and presynaptic PTPσ receptor. The SEC-profile of SALM3-PTPσ complex shows a single major peak, and the frames (350-365) with similar Rg-values, as calculated from the X-ray scattering intensities were selected for further analysis (**Fig. 3a**). The SALM3_LRR-Ig_ and PTPσ_Ig1-Ig3_ structures were modelled based on the known crystal structures, as described in the methods. Rigid body fitting of the complex and *ab initio* envelope modelling were done against the obtained solution scattering data. The complex structure was modelled by rigid body fitting, constraining the LRR dimer structure and the ligand binding interface, while allowing the SALM3 LRR-Ig domain hinge region to rotate. The starting model was based on the SALM5-PTPδ complex crystal structure (PDB 5XNP)^17^. The final rigid body refinement results against the SAXS data demonstrate that a 2:2 complex between SALM3 and PTPσ is formed similar to SALM5, with the noted flexibility at the LRR-Ig domain hinge region of SALM3. Both models with P1 and P2 symmetry fit the data well with Χ^2^-values of 1.7 and 2.5 respectively (**Fig. 3d**,**e**,**f** and Supplementary **Fig. S3**). The results suggest that allowing conformational freedom gives a better fit to the measured data. We have shown previously that the position of the Ig-domain is flexible relative to the LRR domain dimer for both SALM3 and SALM5^19^. This is the case also for the ligand bound structure in solution. The results for the complex structure indicate some flexibility or an equilibrium between different conformation states that can be modelled as presented with the SAXS data, and the 2:2 complex is not necessarily completely fixed and symmetrical in solution or *in vivo*. Furthermore, the *ab initio* calculated envelope can be fitted well with the rigid body models (**Fig. 3g, h**). Consequently, both methods give a result that fit a 2:2 complex best. We further confirmed that 1:1 and 2:1 complexes do not fit to the data (Supplementary **Fig. S3**). Thus, we here show that SALM3 forms a complex with the presynaptic PTPσ receptor with a similar mechanism to SALM5 binding to PTPδ, and that we observe that the structure exhibits certain amount of flexibility that may be requirement for the molecular environment in the synaptic cleft between the two cellular membranes. Based on the SAXS data, we further validated the interaction sites in the SALM3-PTPσ complex by site-directed mutagenesis and cell-based binding and heterologous synapse formation assays.

### Mutational analysis of SALM3-PTPσ interface

The recent SALM5-PTPδ complex crystal structures at 3.7 Å and 4.2 Å resolution showed that the LRRCT-region and Ig-domain of SALM5 interact with the Ig2 and Ig3 domains of PTPδ, and the presence of meB sequence enhanced the interaction^17, 18^. We also reported in our earlier study^19^ using surface plasmon resonance (SPR) that the interaction of SALM3 and PTPσ is enhanced by ten-fold in presence of meB.

Multiple sequence alignment of SALM3 and SALM5 sequences (Supplementary **Fig. S2**), and the amino acid conservation plotted on the SALM3-PTPσ complex model (**Fig. 4a, b, c**) show that the PTPσ interacting residues of SALM3 are mostly conserved in SALM3 and SALM5 in the vertebrates (**Fig. 4a;** Supplementary **Fig. S2, Table S2**). In addition, in the LRR domain dimer structure of SALM3, the concave surface shows large number of conserved residues between SALM3 and SALM5 based on sequence analysis (**Fig. 4a**)^19^. This might indicate a potential binding site, the ligand for which remains to be identified.

**Figure 4.**
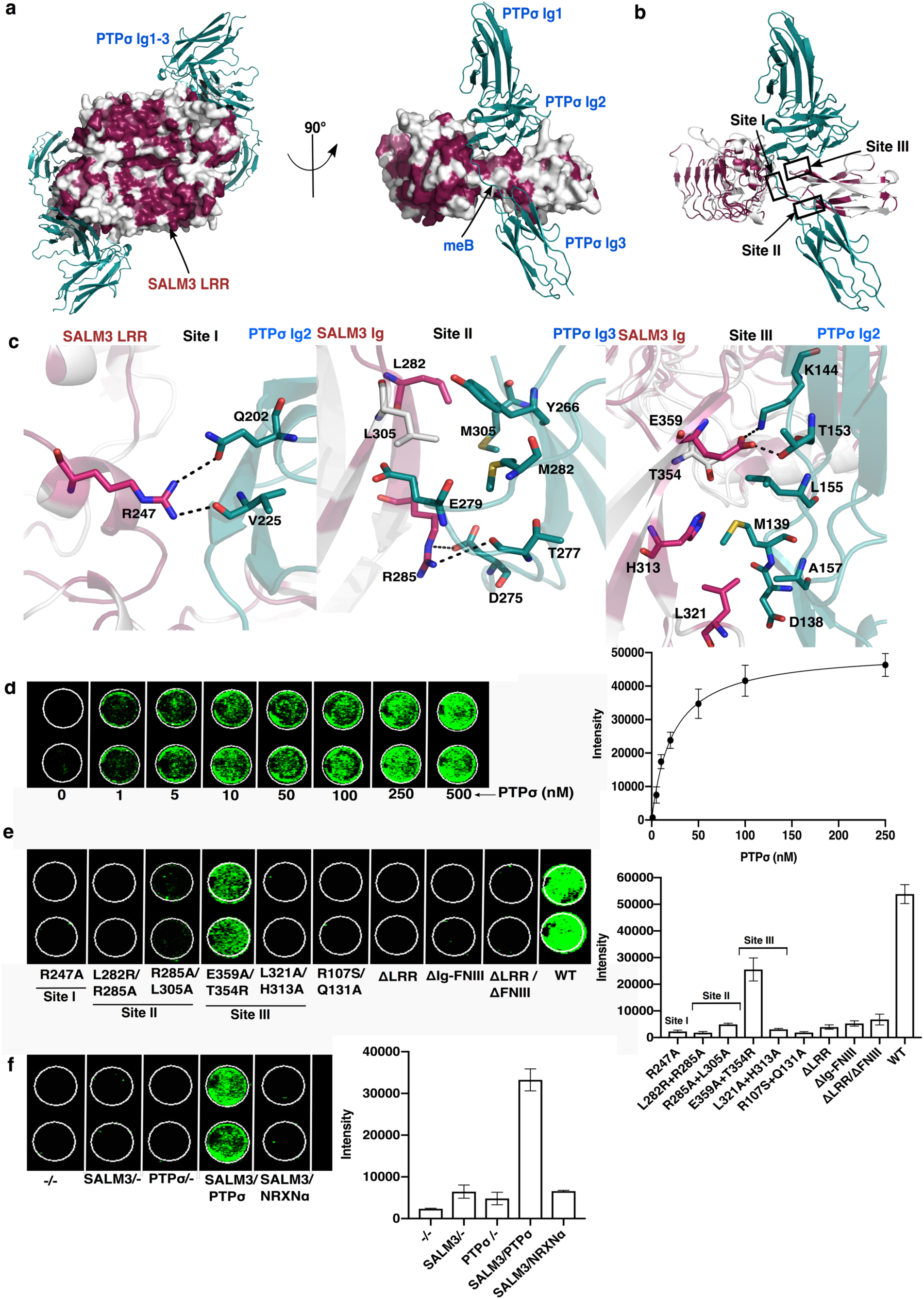
Mutational studies of SALM3-PTPσ interface. a) The SALM3-PTPσ complex model showing the conserved residues in SALM3 LRR-Ig model and the binding interface to PTPσ in the dimeric complex, orthogonal to the SALM3 dimer interface (left), and from the side with PTPσ Ig domains and meB site indicated (right). Red color indicates the most conserved residues in SALM3 LRR-Ig as calculated by CONSURF^43^, PTPσ is colored in teal. b) Interaction sites between SALM3 and PTPσ based on the SALM5-PTPδ crystal structure (PDB ID: 5XNP), are shown as labeled in the figure. c) Close-up view of the predicted interacting sites for the SALM3-PTPσ complex, as labeled in the figure; from the left Site I, Site II and Site III. Residues involved in hydrogen bonding (dashed line) and hydrophobic interactions are shown, colored as in a) and for visual clarity non-carbon atoms color-coded according to atom (nitrogen: blue, Sulphur: yellow, oxygen: red). d) Cell binding assay of soluble PTPσ-Fc (0-250 nM) to SALM3 on the HEK293T cells. Detected in-cell Western blot signals (right) and equilibrium binding curve (left). e) Cell-based binding study of the surface expressed WT SALM3 and mutants to PTPσ on the HEK293T cells. Detected in-cell Western blot signals (right) and binding intensities of PTPσ to WT SALM3 and mutants (left). f) Controls for the cell binding assay. From right; plain (HEK293T) cells (-/-), HEK293T cells with SALM3 transfected (SALM3/-), plain HEK293T cells with ligand PTPσ-Fc added (-/PTPσ), HEK293T (SALM3 expressed) with ligand PTPσ-Fc added (SALM3/PTPσ-Fc) and HEK293T (SALM3 expressed) with ligand NRXN-α-Fc added (SALM3/NRXNα).

Based on the sequence analysis, and in particular the SAXS structure of SALM3-PTPσ complex, we suggest that the binding interface of SALM3-PTPσ is similar to SALM5-PTPδ, with the observed dependence on the presence of meB^14, 19^. In order to verify this, we generated several deletion and point mutations to SALM3 extracellular domain at the SALM3-PTPσ ligand interface modelled based on the binding interface interactions of SALM5-PTPδ structure. Mutants were expressed in HEK293T cells, and tested for interaction with PTPσ-Fc fusion protein using an in-cell western binding assay. Initially, we measured the apparent binding affinity of soluble dimeric PTPσ-Fc to SALM3 on the cell surface to be 22.32 ± 5 nM (**Fig. 4d**). The negative controls showed no detectable signal (**Fig. 4f**), which confirmed the specificity of the assay. To assess the effect of the point mutations, we used 150 nM PTPσ-Fc.

Firstly, SALM3 ΔLRR, SALM3 ΔLRR+ΔFn and SALM3 ΔIg-Fn domain deletion mutants were expressed on the surface of HEK293T cells, but showed no interaction with PTPσ (**Fig. 4e**). The cell-based binding studies on SALM3 deletion mutants verified that both the LRR and Ig-domain of SALM3 are involved and necessary for the PTPσ interaction, and neither alone is enough for binding to the PTPσ. Point mutations on the SALM3 LRR and Ig-domains were analysed for PTPσ interaction to probe and verify more closely the importance of binding interfaces of SALM3-PTPσ complex.

The binding interface in SALM3-PTPσ complex is formed by three interaction sites (**Fig. 4b**, Supplementary **Table S2**), similar to SALM5-PTPδ (PDB 5XNP, 5XWT)^17, 18^. In site I the SALM3 LRRCT forms an interface with the Ig2-domain of PTPσ, in site II SALM5 Ig-domain interacts with Ig3-domain of PTPσ and in site III the SALM3 Ig-domain binds to Ig2-domain of PTPσ^17^. Here, we generated mutations in SALM3 on these three sites (**Fig 4b, c**) in order to validate the SALM3 complex structure.

In the case of SALM3-PTPσ complex, in site I, Arg247 (SALM5 Arg253) forms hydrogen bond with the PTPσ Gln202 and back bone carbonyl of Val225 (**Fig. 4c**). In the cell-based binding assay SALM3 Arg247Ala mutant shows no interaction with PTPσ (**Fig. 4e**). This confirms its critical importance for the interaction for SALM3 with PTPσ. In an earlier study^18^, it was reported that SALM5 Arg253Ala decreased the binding affinity with PTPδ eight-fold compare to the wild-type (WT). The site consists of additional hydrophobic interactions by residues Pro212, Cys 246, Leu249 and Trp250 in SALM5 with PTPδ, corresponding to SALM3 Pro210, Cys240, Leu243 and Trp244 residues, in equivalent positions (Supplementary **Table S2**). Sequence analysis further confirms that the residues are highly conserved in SALM3 and SALM5 (Supplementary **Fig. S2 and Table S2**). Essentially all the PTPσ residues involved are conserved on all three interfaces (Supplementary **Fig. S4**).

In site II, SALM3 Arg285 (SALM5 Arg291) forms an ion pair with PTPσ Asp275, and hydrogen bonds to the main chain carbonyl group on Thr277 of PTPσ (**Fig. 4c**). These interactions are conserved between the two complexes. However, not all interface residues in the SALM Ig-domains are conserved. SALM3 Leu305 (Arg311 in SALM5) binds to a hydrophobic pocket surrounded by Met282, Tyr266 and Glu279 side chains (**Fig. 4c**). This is different in SALM5 where the Arg311 hydrogen bonds to adjacent PTPδ Tyr257 and backbone carbonyl groups at Glu270-Asp271. These amino acid differences at the binding site might lead to slight changes in specificity or affinity *in vivo*.

In addition, at site II in the SALM3 complex Leu282 (SALM5 Leu288) forms hydrophobic interactions with PTPσ Met305 and the peptide main chain at the turn residues 306-307 of the PTPσ Ig3-domain (**Fig. 4c**). The cell-based binding assays of SALM3 Arg285Ala-Leu282Arg and Arg285Ala-Leu305Ala double mutants with PTPσ showed no signal for interaction (**Fig. 4e**), and thus verify the importance of site II and Arg285, Leu282 and Leu305 for PTPσ binding. In previous studies SALM5 Leu288Ala mutation was reported to decrease affinity by 10-fold^18^, and SALM5 Lys309 and Arg311 when mutated to Asn and Thr to introduce an N-glycosylation site in place of Lys309 also abolished binding^17^.

In site III of the model, Glu359 (SALM5 Glu365) forms hydrogen bond with the PTPσ Thr153, and a salt bridge to PTPσ Lys144. SALM3 Leu321 and His313 (SALM5 Leu327 and His319) form hydrophobic interactions with Asp138, Met139 and Ala157 of PTPσ (**Fig. 4c**). SALM3 Thr354 is also part of the interface and packs against Leu155 of PTPσ. An earlier study17 reported that based on Ala-scanning mutagenesis that SALM5 Ser360Ala-Se329Ala double mutant (Ser360Ala equal to SALM3 Thr354) significantly reduced the interaction with LAR^18^. Our cell-based binding study showed that SALM3 Leu321Ala-His313Ala double mutation abolishes the interaction with PTPσ, and SALM3 Thr354Arg-Glu359Ala double mutant decreases binding towards PTPσ by ∼50 % but did not abolish the interaction (**Fig. 4e**). Thus, in site III the hydrophobic interactions by Leu321 and His313 appear to be most critical for the affinity in SALM3 of the residues at this interface **(Fig 4c**,**e)** and Supplementary **Table S2**). Overall, all three sites observed as binding interfaces for SALM5 appear to be functionally equivalent in SALM3, and our mutational data on the interfaces nicely complements the earlier limited mutational studies on SALM5^17, 18^.

Finally, for SALM5, it was reported that the Asn158Ala and Gln134Asn SALM5 mutations (equivalent to SALM3 residues Asn156 and Gln131) that interfere with the LRR dimerization hardly affect the binding to PTPδ^18^. Our cell-based binding studies show that the SALM3 Arg107Ser-Gln131Ala mutations at the dimer interface interfere directly with PTPσ binding (**Fig. 4e**), which is in contrast to SALM5 results. This suggests dimerization on cell surface could be direct pre-requirement for activity, at least in the case of SALM3, and potentially based on the highly similar interaction also in SALM5.

### The effect of SALM3 variants on presynaptic differentiation

Since SALM3 functions as a synapse organizer inducing presynaptic differentiation through binding to presynaptic LAR-RPTPs, we tested the effect of the SALM3 mutations on presynaptic differentiation activity using heterologous synapse-formation assays. SALM3 mutants expressed in HEK293T cells were cultured with hippocampal neurons (DIV9), and the presynaptic differentiation was detected by immunostaining against the presynaptic marker synapsin I (**Fig. 5**). The synaptogenic activity of SALM3 was clearly detected with the WT SALM3, but the interface mutants targeting the SALM3-PTPσ interaction abolished synapsin I clustering (**Fig. 5**). In addition, Thr354Arg-Glu359Ala double mutant at site III showed significantly decreased activity, despite exhibiting 50% residual binding at saturating concentration towards the PTPσ-Fc (**Fig. 4e**). Also the Leu282Arg mutation alone at site II was sufficient to compromise the SALM3 activity in inducing presynaptic differentiation (**Fig. 5**). In case of SALM5, it was previously reported^18^ that SALM5 Arg253Ala (Site I of SALM5-PTPδ complex), SALM5 Leu288Ala (Site II) and SALM5 Ile358Ala (Site III) mutants severely reduced the synaptogenic activity. In addition, the Arg107Ser-Gln131Ala SALM3 mutant that interferes with the SALM3 LRR dimerization also abolished the synaptogenic activity (**Fig. 5**). These results suggest that SALM3 mediated dimerization is critical for interaction with PTPσ and the formation of SALM3-PTPσ *trans*-synaptic signaling complex to induce presynaptic differentiation. Overall, the binding assays results are well correlated with data obtained from heterologous synapse formation analyses. This further suggests that all interactions that are important for the complex formation with PTPσ are also required for the synaptogenic activity. Thus the results verify the biological significance of all the three interaction sites, and the fact that the described SALM3-PTPσ interactions are necessary and sufficient for presynaptic differentiation by SALM3. Recently Xie *et. al*^29^ reported that LAR D1 phosphatase domains can dimerize to promote clustering and activation of synaptic differentiation. We suggest that the dimeric 2:2 complex formation by SALMs on binding to LAR-RPTPs in turn may promote the intracellular homophilic interactions of LAR-RPTPs and further activation of presynaptic differentiation.

**Figure 5.**
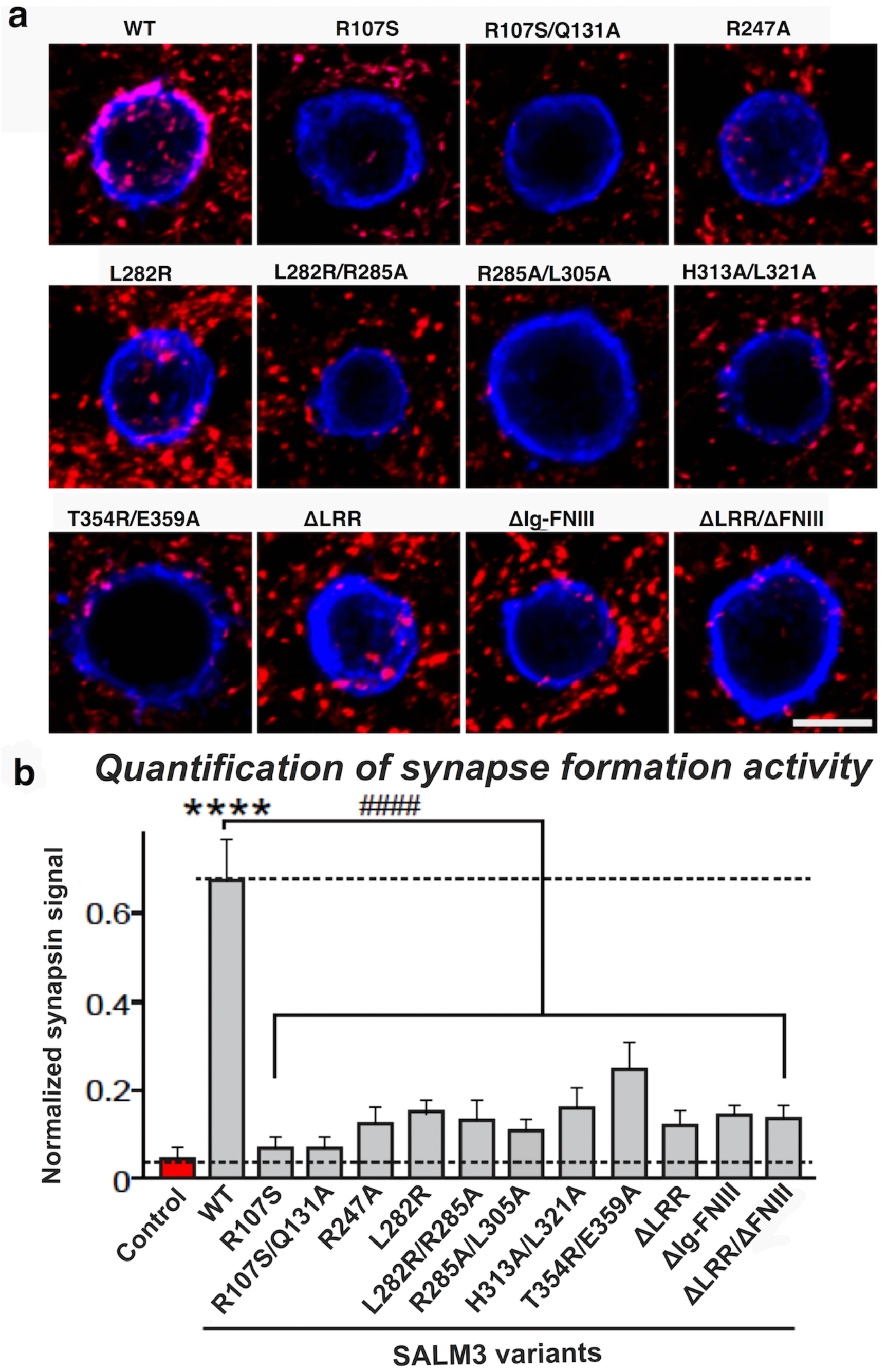
Analysis of SALM3 variants in heterologous synapse formation assay. Representative images of the heterologous synapse-formation activities of SALM3 variants. Cultured Neurons were cocultured from DIV9 to DIV11 with HEK293T cells expressing the indicated WT-SALM3 protein, its point mutants, or deletion variants. Neurons were then stained with antibodies against HA (blue) and synapsin (red). Scale bar = 10 μm (applies to all images). The synapse-forming activity was quantified by measuring the ratio of synapsin staining intensity (red) to HA immunoreactivity intensity (blue). Data are presented as mean ± SEM (\*\*\*\**p* < 0.0001 and ^####^*p* < 0.0001; n = number of neurons as follows: WT, n = 17; R107S, n = 11; R107S/Q131A, n = 13; R247A, n = 17; L282R, n = 16; L282R/R285A, n = 22; R285A/L305A, n = 17; H313A/L321A, n = 16; T354R/E359A, n = 14; ΔLRR, n = 17; ΔIg-FNIII, n = 17; and ΔLRR/ΔFNIII, n = 15). Scale bar, 10 μm (applies to all images).

## Conclusions

SALM3 was the first protein in the SALM-family to be characterized as a ligand for presynaptic LAR-RPTPs^14^. We have now resolved its molecular mechanism of action and mode of oligomerization, which we conclude are similar with SALM5. Taken together, we conclude that dimerization might be a universal mechanism that is conserved across all SALM family proteins and that LAR-RPTP recognition occurs via the same binding interfaces. We hypothesize that SALM3 dimerization promotes presynaptic differentiation by bringing the LAR-RPTP receptors together and enhancing their clustering. Further studies are warranted to systematically test how SALM dimerization is involved in mediation of various synaptic functions and LAR-RPTP activation.

## Materials and methods

### Cloning, protein expression and purification

Mouse SALM3 LRR (residues 17-284) was cloned into Drosophila pRMHA3 expression vector^30^ with CD33 signal sequence at the N-terminus and a C-terminal Fc-tag preceded with a Prescission protease cleavage site. The cDNA for SALM3 was obtained from ImaGenes GmbH. The cloning and protein purification of SALM3 LRR-Ig (residues 17-367) and mouse PTPσ Ig1-3-meB^+^ (resdiues 33-331) have been described in earlier^19^. For cell-based binding assays, the mouse SALM3 LRR-Ig-Fn (residues 17-510) and the deletion mutants SALM3 LRR (residues 17-284) “ΔIg-Fn”, SALM3 Ig (residues 280-367) “ΔLRR+Fn” and SALM3 Ig-Fn (residues 280-502) “ΔLRR” were cloned in the pDisplay plasmid vector. The constructs contained an N-terminal HA-tag followed by the cloned SALM3 sequence, and at the C-terminus Myc-tag followed by transmembrane domain. For mutational studies, we used site directed mutagenesis^31^ to mutate the plasmid constructs of SALM3 LRR-Ig-Fn. The oligonucleotide pairs used for the cloning and mutagenesis are listed in Supplementary **Table S4**.

For SALM3 LRR, the protein expression was verified by transient transfection and western blot detection with goat anti-human polyclonal horse radish peroxidase (HRP) conjugated antibody (Abcam ab98567). A stable *Drosophila* S2 cell line for SALM3 LRR expression was then generated for the protein purification, and protein was expressed in standard manner as described previously^19^. The protein with the C-terminal Fc-fusion was affinity purified with protein-A sepharose (Invitrogen). Samples were eluted with 0.1 M glycine pH 3.0 directly to a neutralizing buffer, with final composition of 60 mM Tris pH 7.4, 300 mM NaCl. The tagged proteins were incubated with Precission protease for 16 h at 4 °C to remove the Fc tag. Precission protease was produced as a GST fusion in *Escherichia coli* BL21 (DE3) from pGEX-6P-1 vector (Addgene). The cleaved Fc-fusion protein were passed through a protein-A column, and flow-through containing the cleaved SALM3-LRR was collected and gel filtered with Superdex 75 10/300 (GE Healthcare) in 60 mM Tris pH 7.5, 300 mM NaCl and concentrated and stored at −80 °C for further use.

### Biophysical characterization

SEC-MALLS was used for the biophysical characterization of size distribution and oligomerization of SALM3 LRR domain construct. The measurements were done at flowrate of 0.5 ml/min over an S-75 Superdex 10/300 column (GE Healthcare) in 20 mM Tris pH 7.4, 150 mM NaCl with a HPLC system (Shimadzu) and a MiniDAWN TREOS light scattering detector, and Optilab rEX refractive index detector (Wyatt Technology Corp.), Sample (100μL) was injected at 1.5 mg/ml. Data were then analyzed with ASTRA 6 software (Wyatt Technology Corp.).

### Crystallization and structure determination and refinement

The SALM3 LRR construct was concentrated to 7.2 mg/ml and exchanged to 20 mM Tris pH 7.4, 100 mM NaCl for crystallization. Thin plate-like crystals appeared from 0.17 M ammonium sulfate pH 7.5, 25% w/v PEG 4000, 10% glycerol at +20 °C. Crystals were harvested directly from the drop and flash frozen for data collection. The crystals diffracted to 2.8 Å and crystallized in space group P2_1_, and were twinned (pseudo-orthorhombic) due to the β–angle being almost equal to 90° **(Table 1)** with high twin-fraction of ca. 34% with the twinning operator h, -k, -l and with two dimers in the asymmetric unit (**Table 1**). The twinning affected the quality of maps and model unavoidably to some extend and complicated the refinement process, however final electron density was of satisfactory quality (Supplementary **Fig. S5**).

The SALM3 crystal structure was solved by molecular replacement with the CCP4 package and PHASER^32^. An initial solution for the SALM3 LRR was found using the LRR domain of SALM5 (PDB 6F20)^19^ as a template. After molecular replacement and initial refinement with REFMAC^33^, the R-factors for the initial LRR domain model were R_work_/R_free_ = 41.2%/46.9%. The model was further refined with PHENIX^34^ and BUSTER^35^ and rebuild with COOT^36^. The final model refined with BUSTER had R-factors of 26.91/28.50% (**Table 1**).

### Small-angle X-ray scattering data collection and analysis

SAXS data on the SALM3 LRR-Ig and PTPσ complex was measured by the SEC-SAXS method at Diamond Lightsource on the B21 beamline. For SALM3 LRR-Ig and PTPσ complex preparation, SALM3 LRR-Ig and PTPσ were mixed in the molar ratio of 1:1.2 of SALM3 LRR-Ig (60 *µ*M) and PTPσ (80 *µ*M) in total volume of 50 *µ*l, followed by incubation at 4 °C for one hour. Sample compartment and exposure cell were cooled to 4 °C. Superdex 200 PC 3.2/30 (GE Healthcare) column was used with flow rate of 0.5 ml/min in 30 mM Tris HCl pH7.5, 150 mM NaCl. Data processing was performed using ScÅtter^37^ and DATASW^38^. Particle shapes at low resolution were reconstructed *ab initio* by DAMMIN^39^ in P1 with P(r) function generated from the data truncated to 0.25 °A^-1^. A total of 10 independent reconstructions were performed, and the models were averaged with the program DAMAVER^40^. Rigid body modeling was performed using CORAL^41^, where available partial crystal structures were treated as rigid bodies while missing residues were modelled as a Cα-chain. The modeling was done with either P1 or P2 symmetry. Porod and dimensionless Kratky plots were calculated for estimation of D_max_ and the overall compactness of the models (Supplementary **Fig. S3**).

### Homology modeling and conservation analysis

Homology models of SALM3 LRR-Ig and PTPσ Ig1-Ig3 structures for SAXS studies and interface analysis were generated based on the known SALM5 (PDB 6F2O) and PTPδ crystal structures (PDB 5XNP) using SWISS-MODEL (https://swissmodel.expasy.org/)^42^. Sequence conservation was analysed and plotted on the SALM3 structure using CONSURF^43^ based on multiple sequence alignment of selected SALM3 and SALM5 LRR-Ig domain amino acid sequences from *Mus musculus, Homo sapiens, Felis catus, Latimeria chalumnae, Danio rerio*, and *Callorhinchus milii* with UniProt/TrEMBL or NCBI sequence IDs Q80XU8, Q6PJG9, M3W7N9, H3BGZ7, B0S5R7 and XP_007909247.1 for SALM3 and Q8BXA0, Q96NI6, A0A337S1F8, H3A0X4, F1Q9H7 and A0A4W3GIX7 for SALM5. The SALM3 LRR-Ig and the PTPσ models were superimposed with the SALM5-PTPδ complex crystal structure (PDB 5XNP), to generate the initial SALM3-PTPσ complex model used as a starting point for SAXS analysis. Multiple sequence alignments were generated with MAFFT^44^.

### Cell-based binding assays

We employed the in-cell western blot method^45^ to detect the SALM3 and PTPσ interaction on cell surface with the Odyssey Infrared Imaging System (LI-COR Biosciences) following manufacturer’s recommendations. In this assay, SALM3 constructs were expressed in HEK293T cells and PTPσ-Fc was added on the cells in a 96-well plate and the binding was detected with anti-human Fc antibody and signal per well was read with the Odyssey Imaging system.

For transfection, HEK293T cells were grown to 90% confluency on a T75 flask using DMEM media with 10% FBS (ThermoFisher) at 37 °C and 5% CO_2_. Cells were seeded into a poly-L lysine-coated 96 well plate at approximately 20 000 cells/well and allowed to grow 24 hours at 37° C until the confluency was 90%. After 24 hours, the cells were transfected with SALM3-pDisplay plasmids (400 *µ*g/well) and polyethylenimine (1.2 *µ*g/well). The transfected cells were incubated at 37 °C for 48 hours. The SALM3 fusion protein expression in transfected cell was detected with Floid™ Cell Imaging system (ThermoFisher) using immunofluorescence.

For immunofluorescence detection, transfected cells were fixed with 4%paraformaldehyde (PFA) for 20 mins and blocked at room temperature with 1xPBS buffer containing 1% BSA for 60 mins, followed by addition of 1:300 dilution primary antibody (rabbit anti-HA, Santa Cruz) and incubation for 30 mins with show shaking at room temperature. The cells were stained with 1:500 dilution secondary antibody (Alexa 488 anti-rabbit, Molecular probes), and the expression was detected with Floid™ Cell Imaging system using green light.

For in-cell western blot experiment, SALM3 transfected HEK293T cells were blocked with EGB buffer^46^ (10 mM HEPES pH 7.2, 168 mM NaCl, 2.6 mM KCl, 2 mM CaCl_2_, 2 mM MgCl_2_, 10 mM D-glucose, and 5% BSA) for 2 hours at room temperature with slow shaking. Blocking buffer was removed and cells were covered in 50 *µ*l of PTPσ-Fc diluted in EGB buffer with 1% BSA. The cells were incubated overnight at 4 °C with gentle shaking. Cells were further treated with 1:200 dilution of secondary antibody (goat anti-human IgG IRDye 800CW), and the SALM3 and PTPσ binding intensity was detected using LI-COR Odyssey (169 *µ*m resolution, medium quality) with channel (800 nm) intensity of 8. For the binding assay a concentration series of PTPσ-Fc (1 to 250 nM) was used, and 150 nM PTPσ-Fc (close to saturation) was used for mutational studies.

### Heterologous synapse-formation assays

HEK293T cells were transfected with the indicated mutants and the WT SALM3 cloned in pDisplay vector (Clontech; listed in **Table S4**). After 48 hours, transfected HEK293T cells were trypsinized, seeded onto cultured hippocampal neurons at DIV9, and then coimmunostained with antibodies against synapsin^47^ at DIV11. Images were acquired by confocal microscopy (LSM700, Carl Zeiss). For quantifications, the contours of transfected HEK293T cells were chosen as the ROI. The fluorescence intensities of synaptic marker puncta, normalized with respect to the area of each HEK293T cell were quantified for both red and green channels using MetaMorph Software (Molecular Devices). All data are expressed as means ± SEM. All experiments were repeated using at least three independent cultures and data were evaluated statistically using a nonparametric Kruskal-Wallis test, followed by Dunn’s multiple-comparison test for *post hoc* groups comparisons (n values used are indicated in the figure legends). Significance was indicated with an asterisk (compared with a value from control group) or hashtag (compared with a value from SALM3 WT-group).

## Acknowledgements

Diffraction data and SAXS data were collected at Diamond light source beam lines I03 and B21. The use of the facilities INSTRUCT-HiLIFE protein crystallization facility (UH) for crystallization and SEC-MALLS measurements is gratefully acknowledged. We are grateful to Maria Vartiainen for kindly providing anti-HA antibody samples for immunofluorescence experiments. Coordinates and structure factors have been submitted with the PDB accession code 6TL8. This work was funded by Jane and Aatos Erkko Foundation (TK), Magnus Ehrnrooth Foundation (SK) and the National Research Foundation of Korea (NRF) funded by the Ministry of Science and ICT (2019R1A2B3B02069324 to JK).

## Author Contributions

SK generated all the constructs, and performed protein expression and purification, cell binding assays, crystallization and other biophysical analyses and contributed to SAXS data analysis and crystal structure refinement, AVS performed the SAXS data analyses, SB performed the heterologous synapse formation assays, JK supervised and designed the heterologous synapse formations assays, TK designed and supervised the overall studies, and contributed to crystallographic data collection and structure solution and refinement. All authors contributed to the writing of the manuscript.

