## Supplementary material for "Structural basis of SALM3 dimerization and synaptic adhesion complex formation with PTPσ"

**Figure S1. Alignment of SALM LRR domains.** Alignment of SALM2 (PDB 5XWU) (red), SALM3 (blue) and SALM5 (PDB 5XNP) (cyan) LRR domains show that the structures are nearly identical. The SALM5 has the long loop connecting 7[th] LRR repeat and LRRCT capping subdomain ordered in the crystal structure.

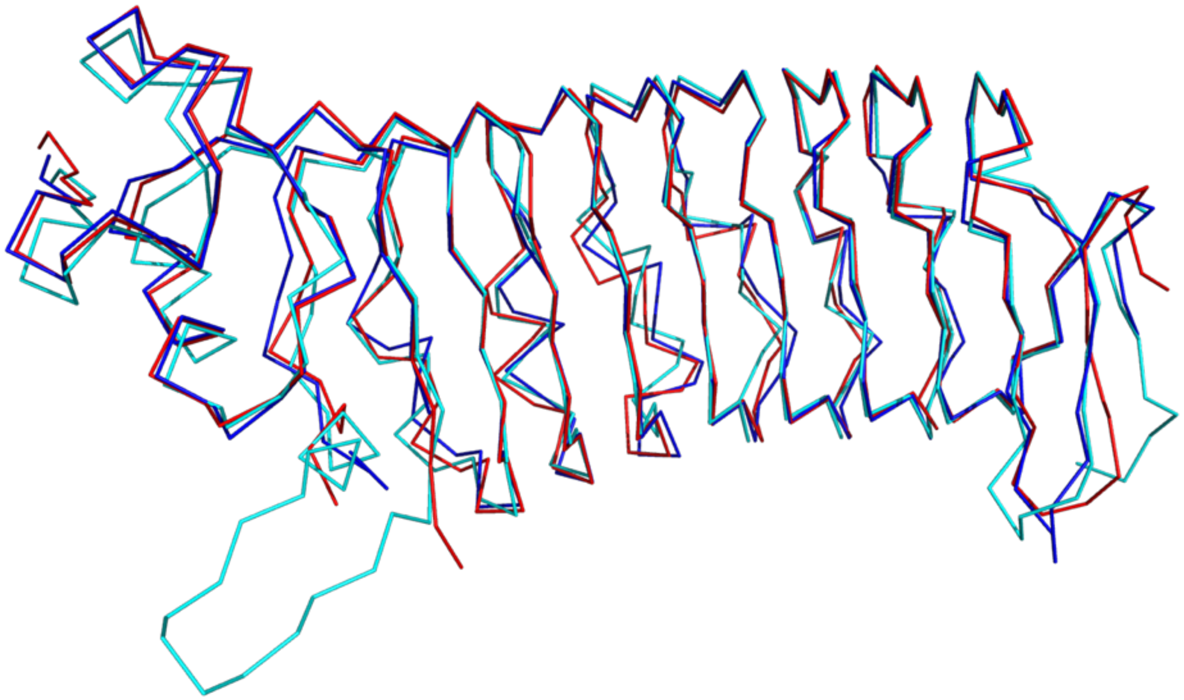

**Figure S2**. **Multiple sequence alignment of SALM3 and SALM5 from selected vertebrate species.** Alignment includes sequences from *Homo sapiens*, Human (Hs), *Felis catus*, cat (Fc), *Latimeria chalumnae*, coelacanth (Lc), *Danio rerio*, zebrafish (Dr) and *Callorhinchus milii*, shark (Cm). On top the secondary structure for the mouse SALM3 structure is shown. Amino acid sequence alignment consists of the SALM LRR and Ig domain regions. The residues located on the SALM3 LRR-dimer interface are labelled with red asterisks (*). The residues involved in SALM5:PTPδ interaction are labelled with black asterisks (*). Conserved residues in SALM3 and SALM5 present in the SALM3 LRR-dimer interface and SALM5-PTPδ interface are highlighted with red color and non-conserved residues with blue color.

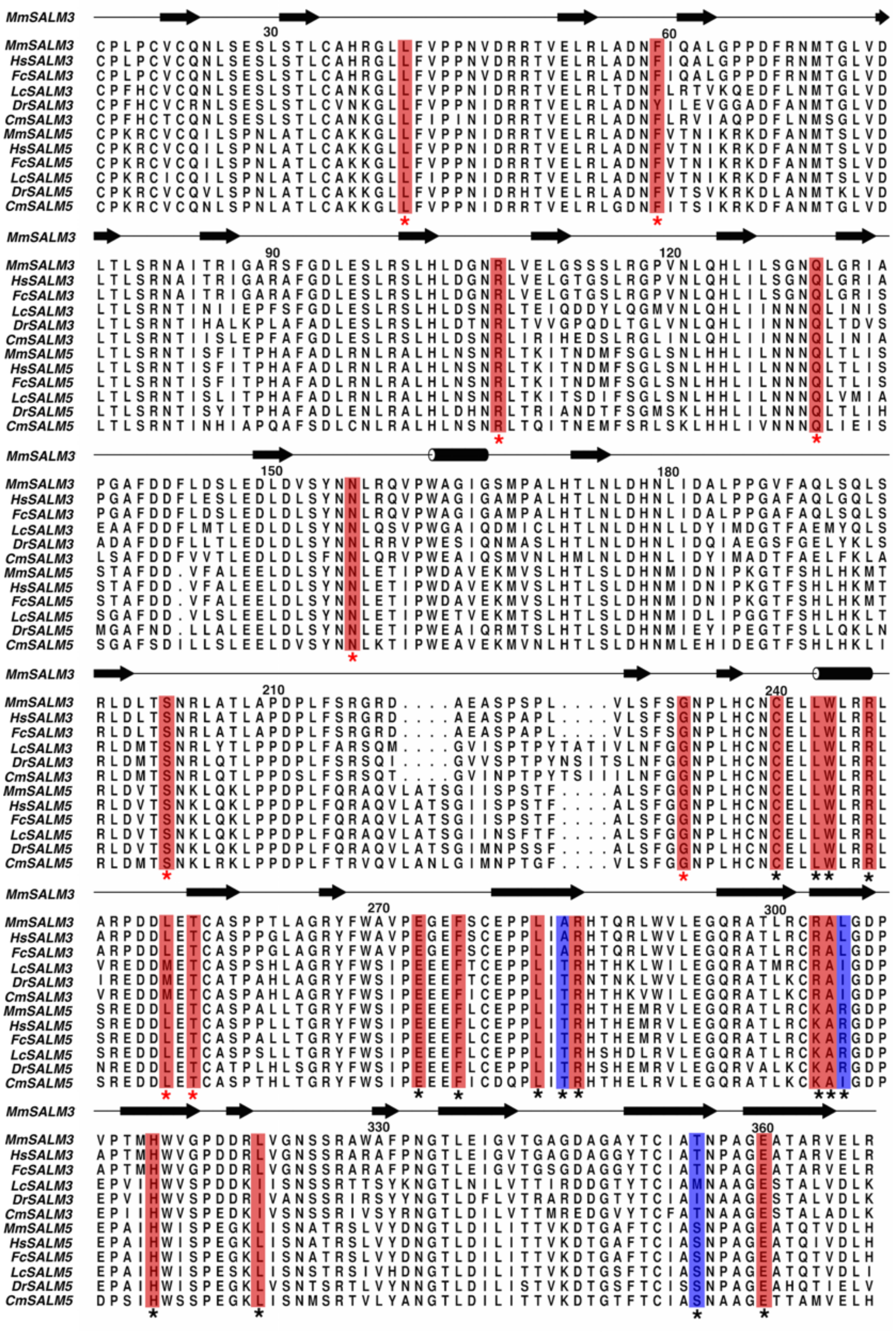

**Figure S3. SAXS analysis of the SALM3-RPTPσ complex.** a) Scattering profiles of SALM3-RPTPσ with the insert plot showing the Guiner region (trend line, blue color) with linear fits. b) Distance distribution function, P(r) of SALM3-RPTPσ complex. c) Dimensionless Kratky plots in comparison with globular BSA (grey) and natively unfolded human hTau40 protein (red). d) Scattering intensities profiles with corresponding fits to SALM3-RPTPσ complex models with different stoichiometry of SALM3 and RPTPσ. Surface structures show LRR domain (red) and RPTPσ (blue). $\chi^2$ values indicate the discrepancy between the experimental data and the scattering intensity calculated from the respective model. Experimental SAXS profiles were appropriately displaced along the logarithmic axis for better visualization and overlaid with corresponding fits.

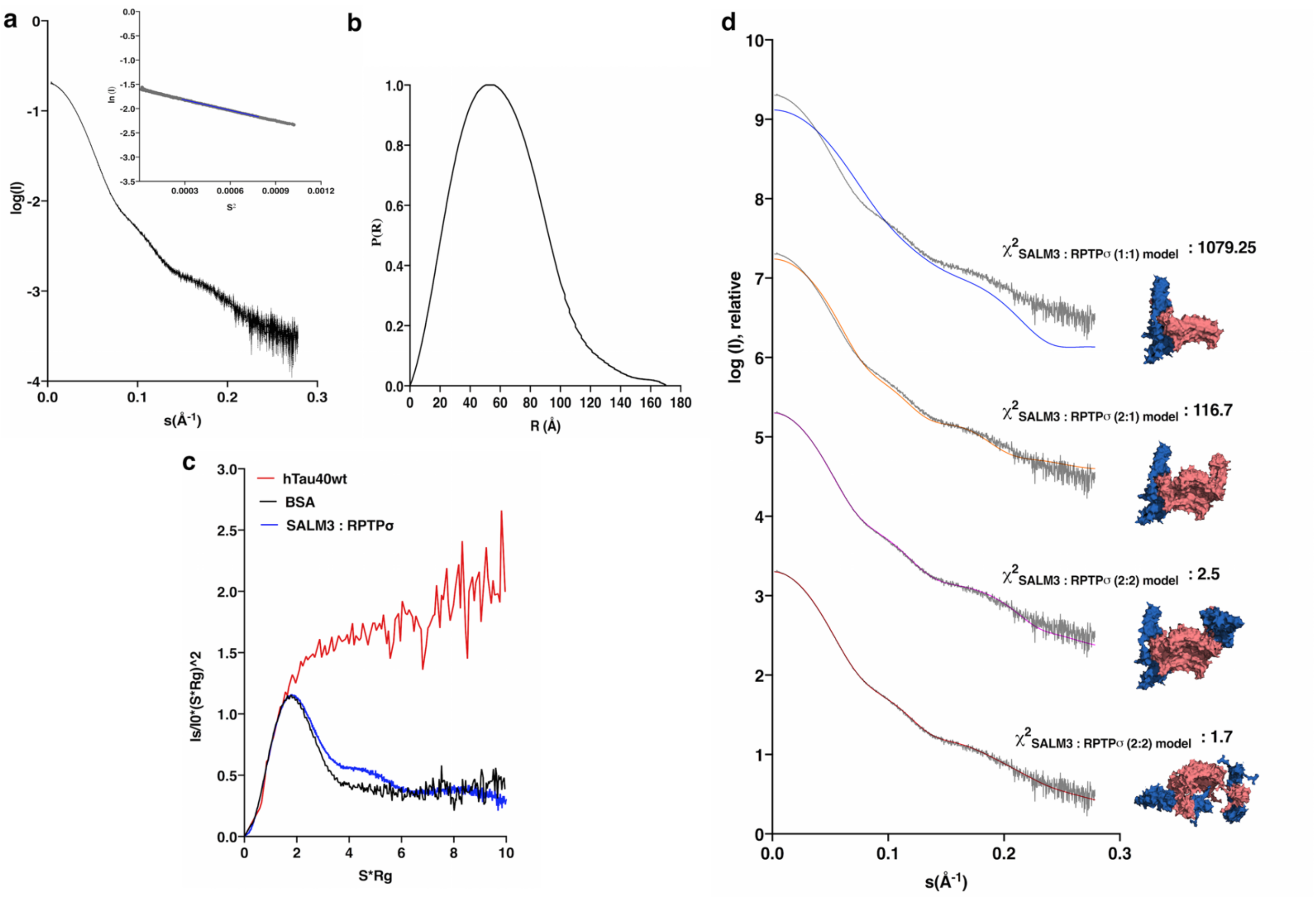

**Figure S4.** Multiple sequence alignment of the three members of human and mouse LAR-RPTPs (LAR, PTPσ and PTPδ). Alignment includes LAR-RPTPs Ig1-3 domains. PTPδ residues involved in SALM5 interaction and conserved in PTPσ are marked. Interface residues identified using PISA server with ≥ 75% buried area percentage. Color coding as in Figure S2.

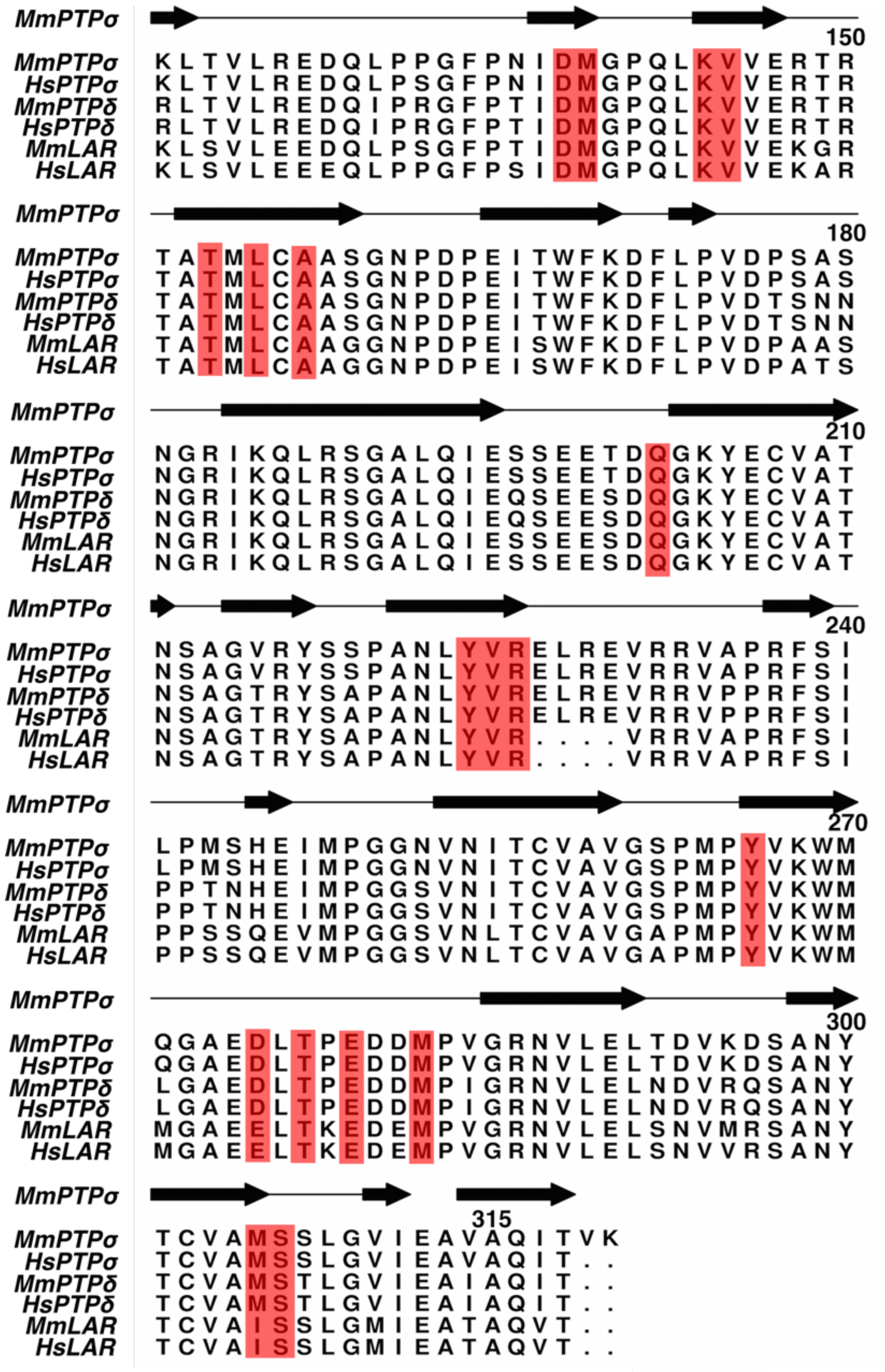

**Figure S5.** Representative $2F_o$-$F_c$ electron density for the SALM3 LRR domain contoured at $1\sigma$ level.

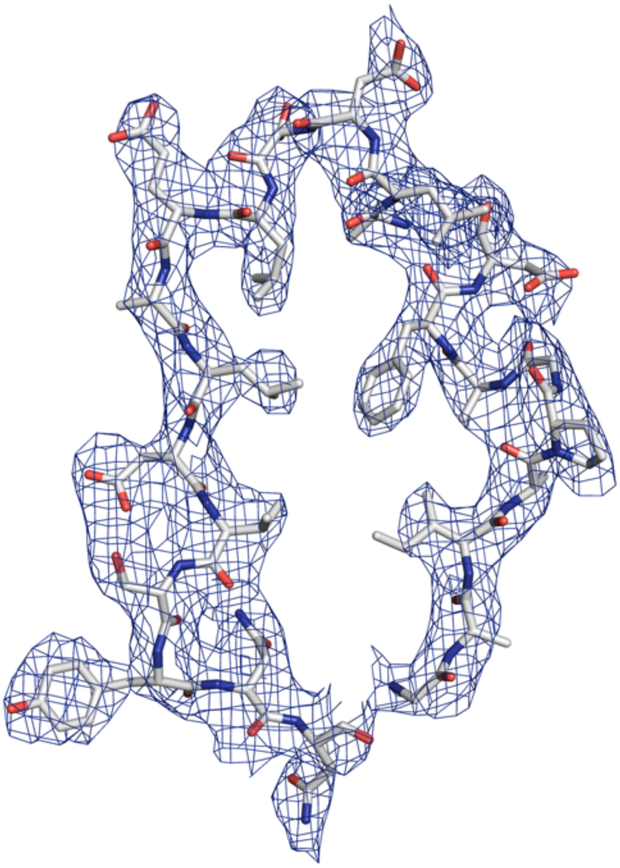

**Table S1.** Conservation of amino acid residues present in SALM3 LRR dimer interface and the corresponding residues in SALM protein sequences. Interface residues in SALM3 structure were identified using the PISA server (https://www.ebi.ac.uk/pdbe/pisa/) with buried area percentage ≥ 75%. Multiple sequence alignment of human SALM proteins was calculated with MAFFT.

| | **SALM5** | **SALM4** | **SALM3** | **SALM2** | **SALM1** |
|---|---|---|---|---|---|
| **Dimer interface** | Leu43 | Leu51 | Leu40 | Leu57 | Leu44 |
| | Phe62 | Phe70 | Phe59 | Phe76 | Phe63 |
| | Arg110 | Arg118 | Arg107 | Arg124 | Arg111 |
| | Gln134 | Gln142 | Gln131 | Gln148 | Gln135 |
| | Asn158 | Asn167 | Asn156 | Asn173 | Asn160 |
| | Ser204 | Ser213 | Ser202 | Ser219 | Ser206 |
| | Gly239 | Gly248 | Gly233 | Gly251 | Gly241 |
| | Leu260 | Leu269 | Leu254 | Leu272 | Leu262 |
| | Thr262 | Ala271 | Thr256 | Thr274 | Thr264 |

**Table S2.** Amino acid residues at SALM3-RPTPσ interface sites and the corresponding residues in the SALM5-RPTPδ complex. Interface residues were identified using PISA server (https://www.ebi.ac.uk/pdbe/pisa/) with buried surface area of ≥ 75%. Interactions in SALM3-RPTPσ interface are indicated according to type of interaction: hydrophobic (black), hydrogen bonds (red) or salt bridges (blue). Differences are marked in bold for SALM5 residues. RPTP residues are all fully conserved (see text for closer description of interactions).

| **Interaction sites** | **SALM** | | | **RPTPs** | | |
|---|---|---|---|---|---|---|
| | **SALM5** | **SALM3** | | **RPTPσ** | **RPTPδ** | |
| Site I | Pro212 | Pro210 | | Leu143 | Leu134 | Hydrophobic |
| | Cys246 | Cys240 | | Val145 | Val136 | Hydrophobic |
| | Leu249 | Leu243 | | Tyr224 | Tyr215 | Hydrophobic |
| | Trp250 | Trp244 | | Gln202 | Gln193 | Hydrogen bond |
| | Arg253 | Arg247 | | Val225 | Val216 | Hydrogen bond |
| Site II | Leu288 | Leu282 | | Met305 | Met296 | Hydrophobic |
| | Ala310 | Ala304 | | Glu279 | Glu270 | Hydrogen bond |
| | Arg291 | Arg285 | | Asp275 | Asp266 | Salt bridge |
| | Arg311 | Leu305 | | Thr277 | Thr268 | Hydrogen bond |
| | | | | Met282 | Met273 | Hydrophobic |
| | | | | Tyr266 | Tyr257 | Hydrophobic |
| Site III | His319 | His313 | | Met139 | Met130 | Hydrophobic |
| | Leu327 | Leu321 | | Asp138 | Asp129 | Hydrophobic |
| | | | | Ala157 | Ala148 | Hydrophobic |
| | Ser360 | Thr354 | | Leu155 | Leu146 | Hydrophobic |
| | Glu365 | Glu359 | | Thr153 | Thr144 | Hydrogen bond |
| | | | | Lys144 | Lys135 | Salt bridge |

**Table S3.** SAXS data collection and structural parameters.

| | **SALM3 LRR-Ig -RPTPσ** | |
|---|---|---|
| **Data collection parameters** | | |
| Beamline<br>Detector<br>Beam geometry<br>Wavelength (Å)<br>*q* range ($Å^{-1}$) *<br>Exposure time (sec)/frame<br>Temperature (K) | B21, DLS<br>PILATUS 1M<br>0.2 x 0.2 mm<br>1.005<br>0.004-0.4<br>2.4<br>298K | |
| Concentration<br>Column | SALM3 (60 $\mu$M) and RPTPσ (80 $\mu$M) in 50 $\mu$l<br>Superdex 200 PC 3.2/30 | |
| **Structural parameters** | | |
| *I(0)* (relative) (from *P(r)*)<br>$R_g$ (Å) (from *P(r)*)<br>*I(0)* (from Guinier)<br>$R_g$ (Å) (from Guinier)<br>$D_{max}$ (Å)<br>Porod volume $V_p$ ($Å^3$)<br>Excluded volume $V_{ex}$ ($Å^3$)<br>(*DAMMIN*/P1) | 0.20± 0.01<br>47.32± 0.01<br>0.20 ±0.01<br>46.41±0.1<br>171.00 ± 10<br>335000 ± 3000<br>400000 ± 4000 | |
| **Molecular mass determination (Da)** | | |
| From Porod volume ($V_p$/1.7)<br>From SAXS MoW2 (Da)<br>From Guinier / *I(0)*<br>From sequence: protein / protein with glycans* (Da) | 197100 ± 1700<br>210000 ± 2100<br>168000<br>149444 / 163400 | |
| **Rigid body Modelling (*CORAL*)** | | |
| Symmetry<br>$\chi^2$ value | P1<br>1.5 | P2<br>2.5 |
| **Modelling parameters** | | |
| Shape reconstruction<br>Symmetry<br>NSD (var) / # of models<br>$\chi^2$ value | DAMMIN<br>P1<br>0.658 (0.016) /10<br>1.327 – 1.344 | |
| **Software employed** | | |
| Primary data reduction<br>Data processing<br>Computation of component model intensities<br>Model representations | *SCÅTTER*<br>*DATASW, PRIMUS*<br>*CRYSOL*<br><br>*PYMOL* | |

* assuming paucimannose-type glycans typical for insect cells

**Table S4**. Oligonucleotide pairs used for cloning and mutation of the gene constructs. On the first line the forward primer, on the second line the reverse primer.

| Plasmid constructs | Plasmid vector | Oligo pairs |
|---|---|---|
| SALM3 $LRR_{17-284}$ | pRMHA3 | 5'-TTTTGAATTCTGCCCGCTACCCTGTGTGTG-3'<br>5'-TTTTGGTACCGGCAATCAGCGGAGGCTCAC-3' |
| SALM3 LRR-Ig-Fn $_{17-510}$ | pDisplay | 5'-ATATATAGATCTTGCCCGCTACCCTGTGTG-3'<br>5'-ATATATGTCGACGGCTGGTAGCGTAGAGAAGTGG-3' |
| SALM3 LRR $_{17-284}$ | pDisplay | 5'-ATATATAGATCTTGCCCGCTACCCTGTGTG-3'<br>5'-ATATATGTCGACAGGCTCACAGGAGAACTCTCCCTC-3' |
| SALM3 $Ig_{280-367}$ | pDisplay | 5'-ATATATAGATCTGAGCCTCCGCTGATTGCC-3'<br>5'-ATATATGTCGACCCGGAGCTCCACTCGGG-3' |
| SALM3 $Ig\text{-}Fn_{280-502}$ | pDisplay | 5'-ATATATAGATCTGAGCCTCCGCTGATTGCC-3'<br>5'- ATATATGTCGACGGCTGGTAGCGTAGAGAAGTGG-3' |
| SALM3 Q131A | pDisplay | 5'-CTCAGTGGCAATGCACTGGGCCGCATCGC-3'<br>5'-GCGATGCGGCCCAGTGCATTGCCACTGAG-3' |
| SALM3 R247A | pDisplay | 5'-CTGCTGTGGCTGCGGGCGCTGGCCCGGC-3'<br>5'-GCCGGGCCAGCCGCGCCAGCCACAGCAG-3' |
| SALM3 L282R | pDisplay | 5'-CTGTGAGCCTCCGCGGATTGCCCGGCACAC-3'<br>5'-GTGTGCCGGGCAATCCGCGGAGGCTCACAG-3' |
| SALM3 R285A | pDisplay | 5'-CTCCGCTGATTGCCGCGCACACACAGCGCCTG-3'<br>5'-CAGGCGCTGTGTGTGCGCGGCAATCAGCGGAG-3' |
| SALM3 R303A/L305A | pDisplay | 5'-CACCCTACGGTGCGCGGCCGCTGGTGACCCTGTACC-3'<br>5'-GGTACAGGGTCACCAGCGGCCGCGCACCGTAGGGTG-3' |
| SALM3 H313A | pDisplay | 5'-GTACCTACCATGGCCTGGGTTGGCCCTG-3'<br>5'-CAGGGCCAACCCAGGCCATGGTAGGTAC-3' |
| SALM3 L321A | pDisplay | 5'-CCTGATGACAGGGCGGTTGGCAACTCTTC-3'<br>5'-GAAGAGTTGCCAACCGCCCTGTCATCAGG-3' |
| SALM3 T354R | pDisplay | 5'-CCTGCATTGCCCGCAACCCTGCTGGTG-3'<br>5'-CACCAGCAGGGTTGCGGGCAATGCAGG-3' |
| SALM3 E359A | pDisplay | 5'-CAACCCTGCTGGTGCGGCCACAGCCCGAGTG-3'<br>5'-CACTCGGGCTGTGGCCGCACCAGCAGGGTTG-3' |
